# Metabolic cofactors act as initiating substrates for primase and affect replication primer processing

**DOI:** 10.1101/741967

**Authors:** Christina Julius, Yulia Yuzenkova

## Abstract

Recently a new, non-canonical type of 5’-RNA capping with cellular metabolic cofactors was discovered in bacteria and eukaryotes. This type of capping is performed by RNA polymerases, the main enzymes of transcription, which initiate RNA synthesis with cofactors. Here we show that primase, the enzyme of replication which primes synthesis of DNA by making short RNA primers, initiates synthesis of replication primers using the number of metabolic cofactors. Primase DnaG of *E. coli* starts synthesis of RNA with cofactors NAD^+^/NADH, FAD and DP-CoA *in vitro*. This activity does not affect primase specificity of initiation. ppGpp, the global starvation response regulator, strongly inhibits the non-canonical initiation by DnaG. Amino acid residues of a “basic ridge” define the binding determinant of cofactors to DnaG. Likewise, the human primase catalytic subunit P49 can use modified substrate m^7^GTP for synthesis initiation.

For correct genome duplication, the RNA primer needs to be removed and Okazaki fragments ligated. We show that the efficiency of primer processing by DNA polymerase I is strongly affected by cofactors on the 5’-end of RNA. Overall our results suggest that cofactors at the 5’ position of the primer influence regulation of initiation and Okazaki fragments processing.

**Visual abstract:** **A. Non-canonical capping of RNA by RNA polymerase.** RNA polymerase uses cellular cofactor as initiating substrate for RNA synthesis, instead of NTP. Then RNA chain grows, while cofactor remains attached and serves as cap. **B. Proposed mechanism of non-canonical initiation of RNA primer synthesis by DnaG primase during replication.** DnaG primase initiates synthesis of the primer for DNA replication using cellular cofactor. Primer stays annealed with the DNA template. DNApolI encounters cofactor, which affects the removal of primer.

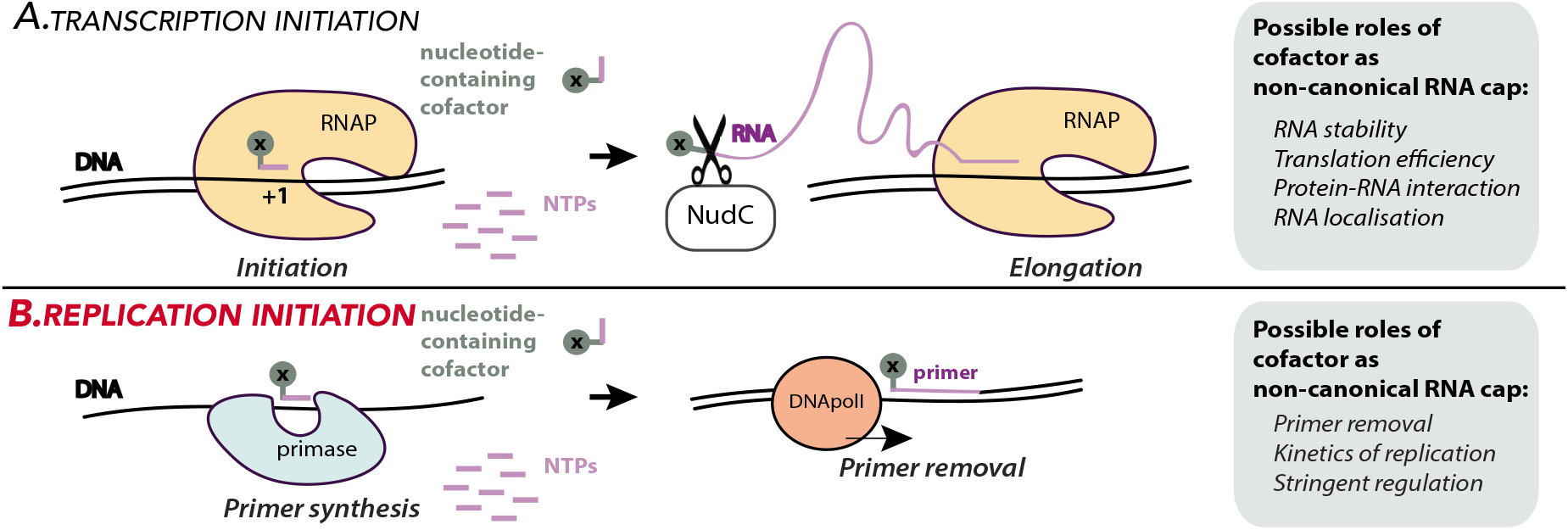

## Introduction

This work was inspired by the recent discovery of the non-canonical RNA capping phenomenon. Many RNA species in bacteria and eukaryotes bear metabolic adenine-containing cofactors at their 5’-end; NAD^+^ (nicotinamide adenine dinucleotide) and DP-CoA (dephospho-coenzyme A) and FAD (flavin adenine dinucleotide), as well as cell wall precursors UDP-GlcNAc (uridine 5’-diphospho-N-acetylglucosamine) and UDP-Glc (uridine 5’-diphosphate glucose) (1, 2). Unlike the classic cap m^7^G, non-canonical caps are installed by the main enzyme of transcription, RNA polymerase (RNAP)(3). It happens during initiation of transcription in a template-dependent manner – ADP-containing cofactors are incorporated at promoters with +1A start sites, i.e. promoters dictating ATP as the initiating substrate (3, 4), UDP-containing cell wall precursors – on +1U promoters (3).

By analogy with the classic cap, there are decapping enzymes for non-canonical caps, in *E. coli* it is NudC (NADH pyrophosphohydrolase of the NUDIX family). NudC processes NADylated RNAs into a monophosphorylated species that are quickly degraded in the cell (5).

Overall it appears that unrelated multi-subunit eukaryotic and bacterial RNAPs as well as the single-subunit RNAPs of mitochondria and viruses can utilize non-canonical initiating substrates (NCISs) and perform RNA capping (6). Another DNA-dependent enzyme initiating *de novo* synthesis of RNA is primase (DnaG in bacteria) which makes primers for replication and present in all organisms. It is not structurally related to either single subunit e.g. mitochondrial RNAP or multi subunit bacteria or eukaryotic RNAPs. In *E. coli*, DnaG recognises a consensus GTC motif and makes a 10-12 nucleotides long RNA primer. Primase acts in concert with other replication proteins. Primase requires DnaB helicase to start synthesis on double-stranded DNA (7). Primase is then displaced and the primer is elongated by the DNA polymerase III to produce the RNA/DNA polynucleotide of a leading strand or an Okazaki fragment of a lagging strand (8). RNA primers need to be removed via the combined actions of DNA polymerase I (PolI) and/or RNaseH before the Okazaki fragments and leading strand are ligated to complete genome replication. Thus, synthesis of an RNA primer by primase is believed to be a rate limiting step of replication (9), tightly coupled to other steps of the replication process. Primase plays a key role during assembly of the replisome (10), regulation of replication elongation and Okazaki fragments length (11, 12) in both bacterial and eukaryotic systems. In bacteria, DnaG primase plays a part in a global concerted response to starvation via starvation alarmone ppGpp which inhibits primer synthesis in nutrient limited conditions (13).

Here we show that *E. coli* primase DnaG starts synthesis of a replication primer using a number of ADP-containing cofactors *in vitro*, including NAD+/ NADH, FAD and DP-coA. This reaction requires amino acid residues of the DnaG “basic ridge” region, and is inhibited by global starvation alarmone ppGpp. We also show that cofactors on the 5’-end of RNA specifically and differentially affect processing of this RNA by DNA polI. Our data suggest that 5’-cofactors influence initiation efficiency and the rate of processing of replication intermediates.

## Results

### *E. coli* DnaG primase initiates RNA primer synthesis using NAD+, NADH, FAD and DP-coA

DnaG primase functions as a low-processive RNA polymerase able to start *de novo* RNA synthesis on a DNA. We wanted to test if primase can initiate synthesis using ADP-containing metabolic cofactors, (structures on Fig. 1A), by analogy with other polymerases.

**Figure 1.**
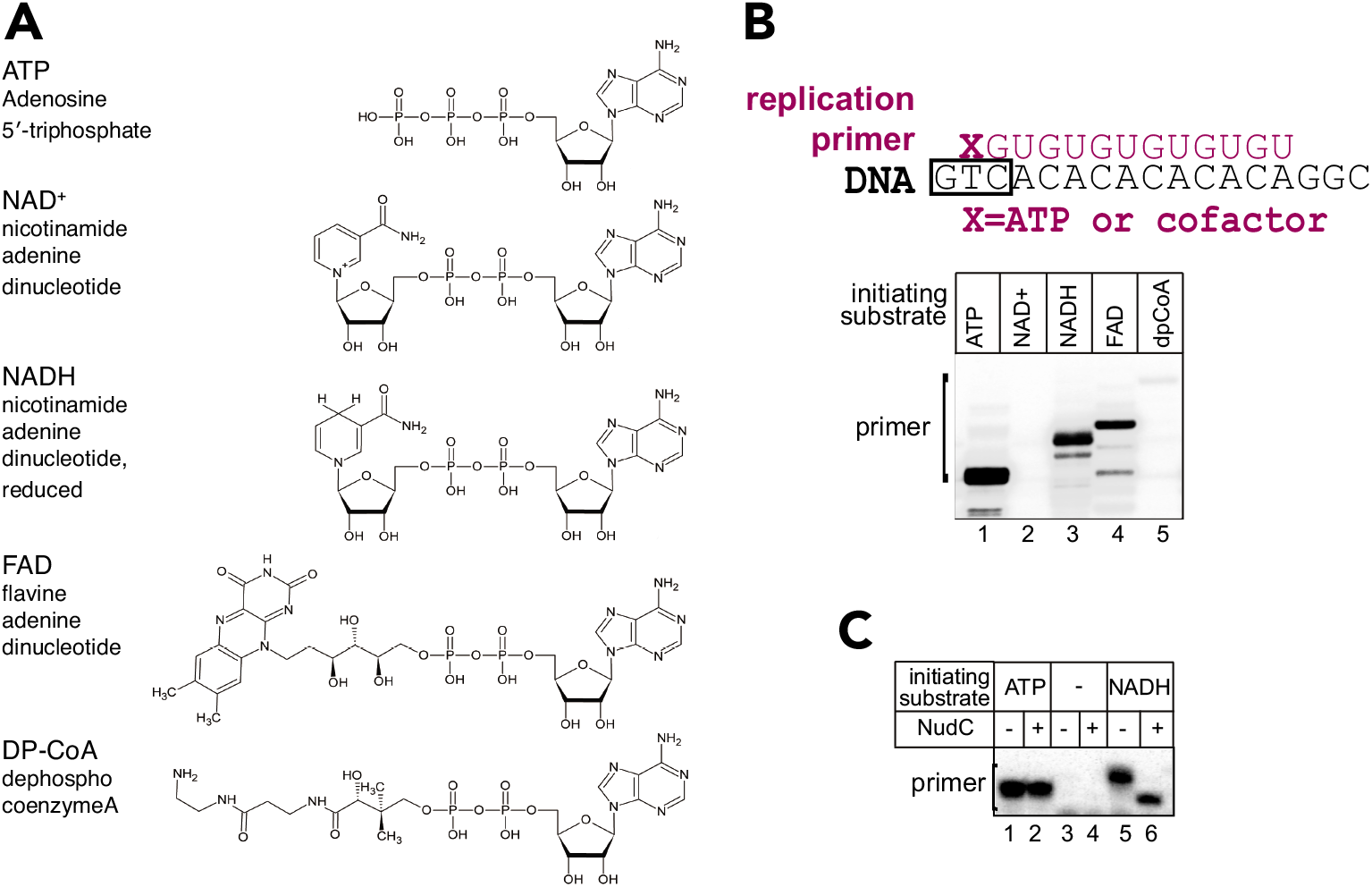
Primase DnaG initiates replication primer synthesis with NAD+, NADH, FAD and DP-coA. **A.** Structures of cofactors molecules (NAD+, NADH, FAD and DP-coA) in comparison to ATP, the preferred initiating substrate of primase. **B.** Replication primer synthesis on single-stranded DNA template, scheme above, with ATP, NAD+, NADH, FAD, and DP-coA as initiation substrates. **C.** RNA primer with 5’-NADH is susceptible for cleavage by NudC nuclease after DnaG was removed from the complex with high salt wash. Note that absence of ATP from the reaction results in an increased amount of non-specific product, which is not susceptible to NudC in lanes 5, 6. We assume that this band results from initiation with GTP present in the reaction.

We used a general priming system, i.e. minimal system in the absence of singlestrand DNA-binding protein. This set-up requires only DnaG for RNA primer synthesis on the short single-stranded DNA template containing GTC recognition motif (scheme on Fig. 1B)(7).

We found that DnaG makes a 13nt long RNA product in a subset of NTPs using either ATP, NAD+, NADH, FAD and DP-coA (Fig. 1B). To avoid confusion and for simplicity, we will refer to the length of RNA with conventional substrates even though cofactors are dinucleotides. Notably, DnaG incorporates NAD+ much less efficiently than NADH, in contrast to other RNAPs, which do not discriminate between NAD+ and NADH (4). We found that affinity to the non-canonical initiation substrates is comparable with their physiological concentrations. Michaelis constants we measured for ATP, NADH and FAD as initiating substrates, were 46.6, 109 and 390 μM, correspondingly. In actively growing in rich media *E. coli* cells, concentrations of ATP, NADH and FAD are 9.6 mM, 100-1000 μM and 200 μM, correspondingly (14, 15).

NADH capped primers were not susceptible to decapping by NudC, unless primase was washed away with high salt buffer, like in the experiment shown on Figure 1C. This result is in agreement with the view that full length primer stays bound to the DNA template and in complex with primase (12), in our case even if the primer contains extra moiety on its 5’-end.

Interestingly, after primase was removed, 5’-NADH of “naked” RNA primer annealed to DNA was susceptible to NudC, in contrast to the published result with 5’-NADH of RNA annealed to RNA (5). Therefore, NudC may have a limited window of opportunity to remove the cap from the primer during active replication and participate in processing of replication intermediates.

### Cofactors substrates do not affect DnaG specificity of initiation

Cofactors are dinucleotides, and therefore can make additional contacts with DNA template at −1 position relative to the start site (scheme on Fig. 2). These contacts may affect initiation specificity by DnaG which recognises GTC motif on a template strand. The central T codes for A and G is a template −1 base. To test if cofactors incorporation depends on the nature of −1 base, we tested synthesis of RNA13 on templates with −1 changed to three other bases. This experiment we performed with initiation substrates concentrations roughly in the region of their corresponding K_m_s (50 μM for ATP and 500 μM for cofactors). We found that primase preferred purines in this position, and the least preferred base is C (Fig. 2). Initiation with cofactors did not change these preferences, suggesting that cofactors do not make specific contacts with −1 base of the template. This result also implies that cofactors as substrates do not change specificity of DnaG initiation and do not lead to spurious initiation.

**Figure 2.**
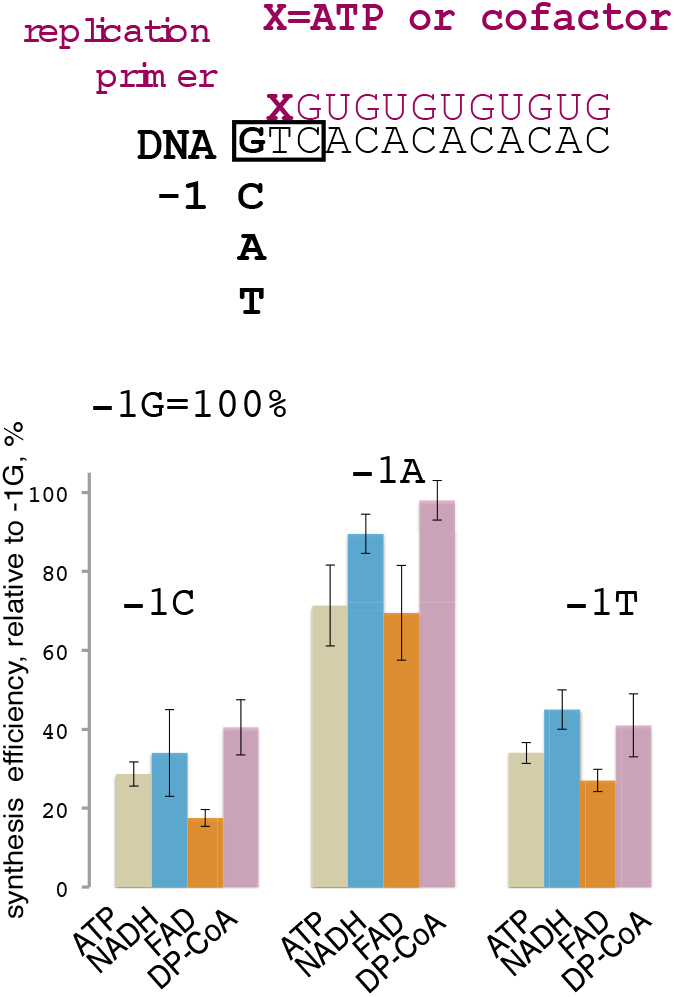
Cofactors do not make specific contacts with −1 DNA template base. Synthesis efficiency of RNA13 started with ATP or NADH, FAD or DP-CoA on DNA templates with C, A or T at position −1 was compared to consensus −1G template. Relative efficiency of synthesis is shown in percentage from −1G template, error bars reflect standard deviations from three independent experiments.

### Initiation of RNA synthesis with cofactors requires basic ridge amino acid residues of DnaG

Since cofactors do not make strong contacts with DNA, they most probably contact DnaG protein itself. A number of amino acid residues, including several in a “basic ridge”, were implicated in initiation nucleotide binding based on sequence conservation amongst primases and structural information for the *Staphylococcus aureus* primase (13). We tested synthesis of a primer by DnaG with amino acid substitutions, K229A, Y230A K241A (basic ridge) and D309A (participating in metal chelation), which were all previously shown to influence initiating substrate incorporation (13, 16), in the presence of DnaB. We found that “basic ridge” substitutions K229A, Y230A, and to some extent K241A specifically inhibited capping with NADH and FAD (Fig. 3), in contrast with D309A. We assumed that NMN and FMN moieties of the corresponding cofactors might make contacts with these amino acid residues either during binding of the initiating substrates or during the very first step of RNA synthesis.

**Figure 3.**
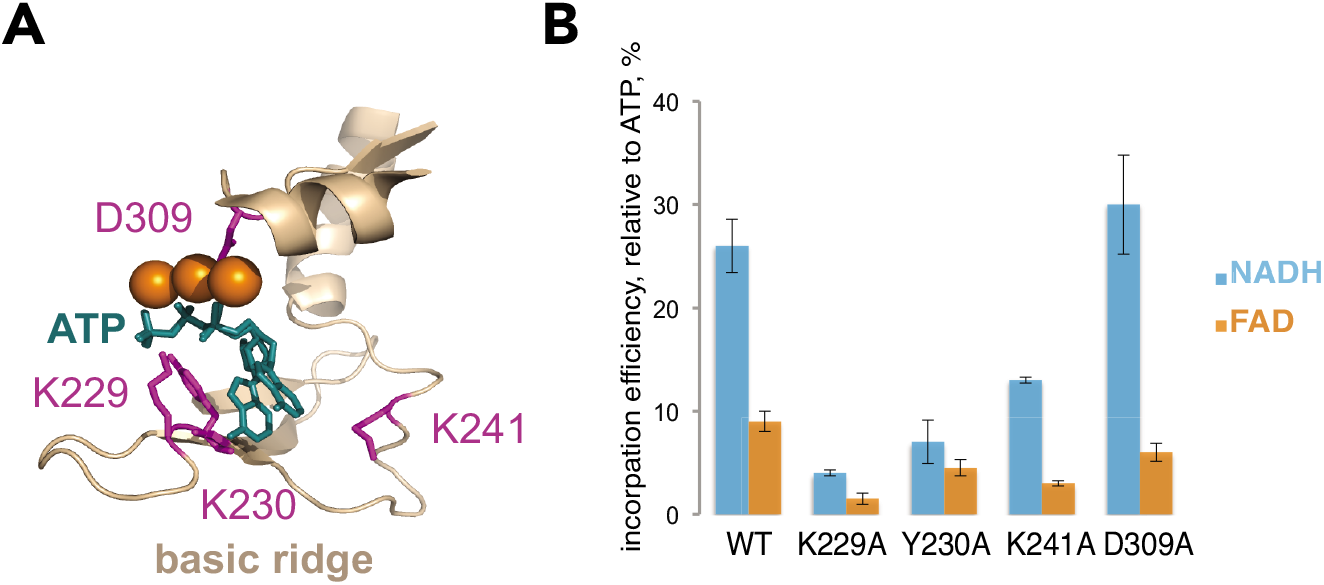
DnaG basic ridge residues affect initiation with NADH and FAD. **A.** Structure of the *S. aureus* DnaG-ATP complex in the vicinity of a catalytic site PDB 4EDG. ATP (in two conformations) is in teal, amino acid residues, which we tested for efficiency of cofactor initiation are in magenta. **B.** Relative efficiency of NADH and FAD utilisation as initiation substrates in comparison to that of ATP, for WT DnaG and mutants with amino acid residues substitutions to alanines at positions 229, 230, 241 and 309. Error bars reflect standard deviations from three independent experiments.

### ppGpp strongly inhibits non-canonical initiation by DnaG

Under nutrient deficient conditions, replication is inhibited via the action of global stringent response regulator, alarmone ppGpp, on DnaG. PpGpp binds DnaG at the active site (13), presumably overlapping with the binding site where caps bind during the initiation of primer synthesis. To test if ppGpp competes with capping cofactors NADH and FAD during initiation we measured the maximal rate of RNA product formation in the presence of increasing concentrations of ppGpp (control reactions were done with ATP as initiator). From the plot of residual activities vs ppGpp concentration, it appears that ppGpp competes with cofactors more efficiently than with ATP (Fig. 4). This tendency correlates with Michaelis constants for the initiation substrates which increase in a sequence ATP-NADH-FAD. Therefore, in the absence of structural information on DnaG complexes with cofactors, we can only suggest that binding site is shared by ppGpp and cofactors. During the stringent response, ppGpp concentration might raise above 150 μM in *E. coli* (17) which is roughly, judging from plot, above or close to ppGpp inhibition constants for synthesis reaction of FAD-RNA and NADH-RNA. This result might additionally mean that *in vivo* replication initiation with cofactors would be inhibited earlier during transition into stationary phase or nutritional downshift.

**Figure 4.**
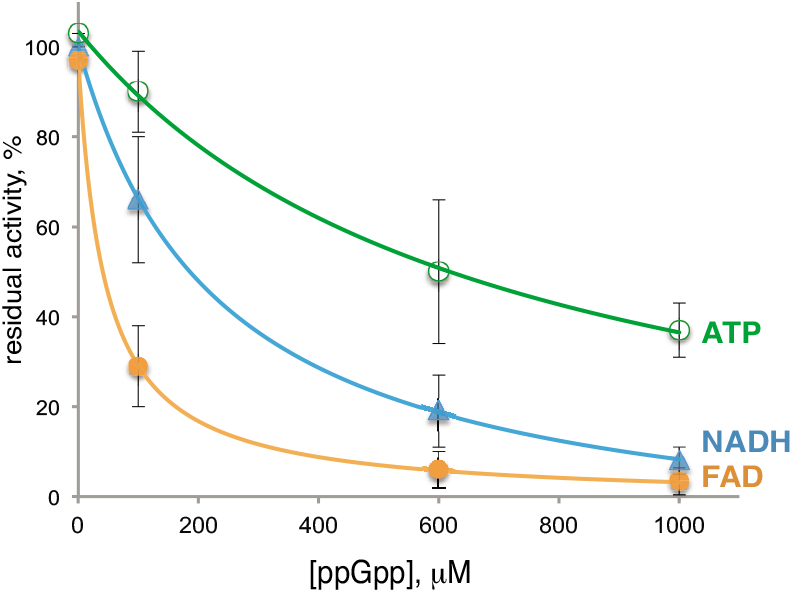
Initiation with cofactors is more susceptible to inhibition by ppGpp than initiation with ATP. Residual DnaG activities are plotted against an increasing concentrations of ppGpp, in percentages from amounts in the absence of ppGpp. Error bars reflect standard deviations obtained from three independent experiments.

### Cofactor 5’-caps differentially affect primer processing by PolI

To complete replication, the leading strand and Okazaki fragments of a lagging strand need to be processed and ligated. The processing involves RNA primer degradation coupled to extension of previous Okazaki fragment by PolI, which possesses 5’-exonuclease and 3’-DNA polymerase activities. We examined if the presence of cofactor on the 5’-end of a RNA primer affects its removal by PolI. This experiment was done using a DNA-RNA construct mimicking part of replication intermediate (Fig. 5, top). The construct consisted of hairpin-containing DNA template with RNA primer (RNA12) produced by DnaG annealed to the single stranded DNA part. DnaG primase was subsequently removed by washing with high salt containing buffer. The DNA-RNA construct was immobilised on streptavidin beads via biotin on DNA, which ensured that the processing observed happens on an RNA annealed to a DNA (See Methods section for more details). Addition of PolI and dNTPs led to a stepwise degradation of RNA, as seen from kinetics of RNA12 degradation on a gel image and the plot below it on Fig. 5. PolI activity was stimulated by the 5’-end NADH; from Fig. 5 it can be seen that full length NADH-RNA12 disappeared in 30 seconds. In contrast, presence of 5’-FAD on the RNA12 greatly reduced the rate of degradation, and FAD-RNA12 was still visible after 4 minutes. ATP-RNA12 was degraded with somewhat intermediate rate. Our data suggest that 5’ cofactors on the primer may influence its removal *in vivo*.

**Figure 5.**
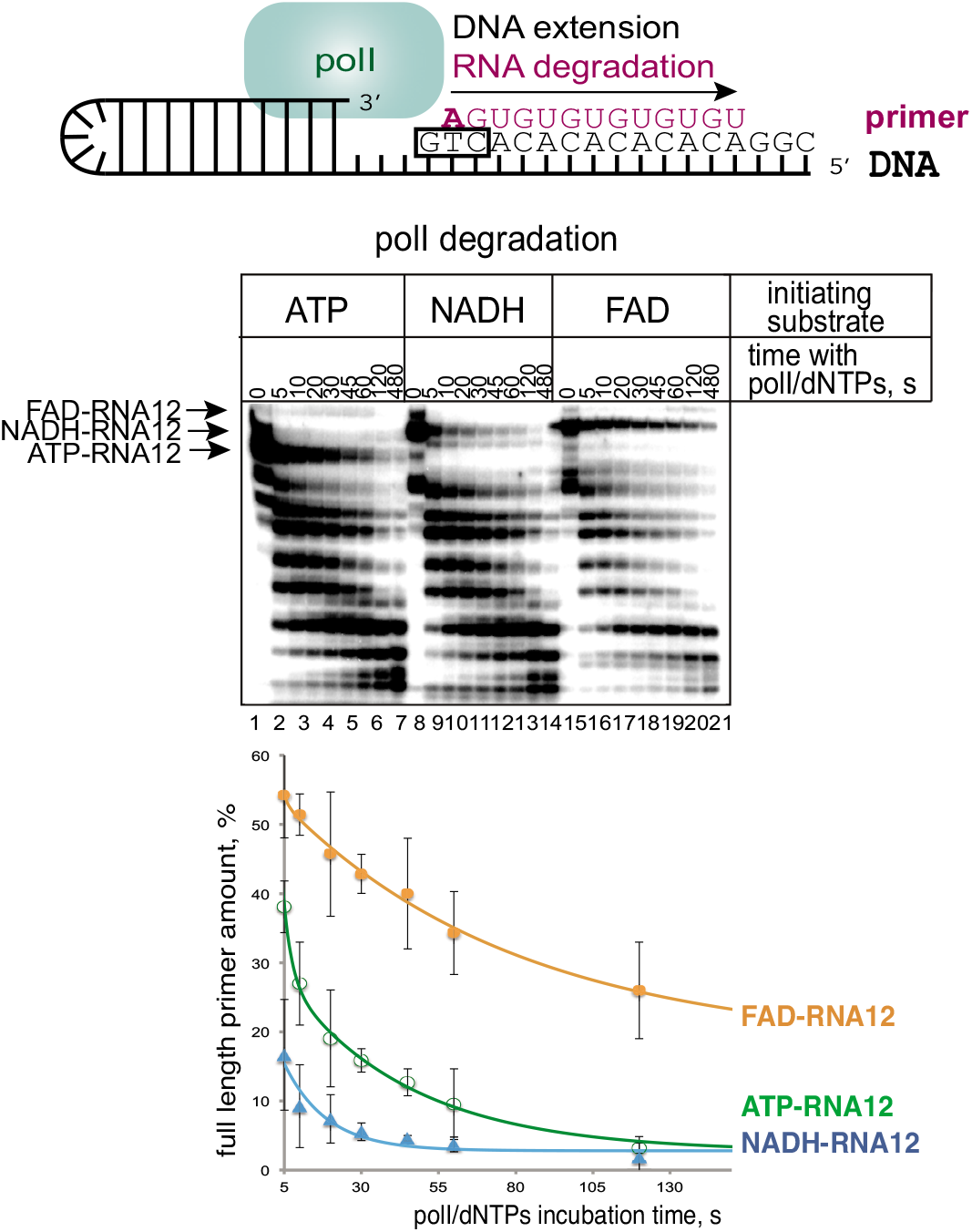
Cofactors at the 5’-end of a primer differentially affect its processing by PolI – NADH speeds it up, FAD inhibits. Scheme of the hairpin DNA substrate with annealed RNA made by DnaG is shown above the gel. On the gel products of time dependent degradation of the ATP-RNA12, NADH-RNA12 FAD-RNA12 made with either ATP, NADH or FAD as initiating substrates is shown. Below the gel image, the same degradation kinetics are presented on a plot. Plot shows amount of the initial full-length of primer as a function of incubation time with PolI and dNTPs. Error bars reflect standard deviations obtained from three independent experiments.

### Eukaryotic primase catalytic subunit P49 uses modified initiating substrate

In addition to prokaryotic system, we wanted to test if the human primase catalytic subunit P49 utilises ADP containing cofactors. We analysed the formation of the first dinucleotide product by this enzyme on the single stranded DNA template (sequence on Fig. 6). We were unable to make this enzyme to start synthesis with ATP at a specific start site, and initiation with GTP was very inefficient on this template (lane 2). Yet, P49 efficiently produced a dinucleotide using m^7^GTP as initiating and UTP as the substrate for the second position (lane 4). This ready incorporation of a modified substrate hints at P49 general low fidelity and propensity to non-canonical initiation. Therefore, our result suggests that P49 might utilise a variety of non-canonical substrates with possible consequences for initiation kinetics and elongation to full length primer (18).

**Figure 6.**
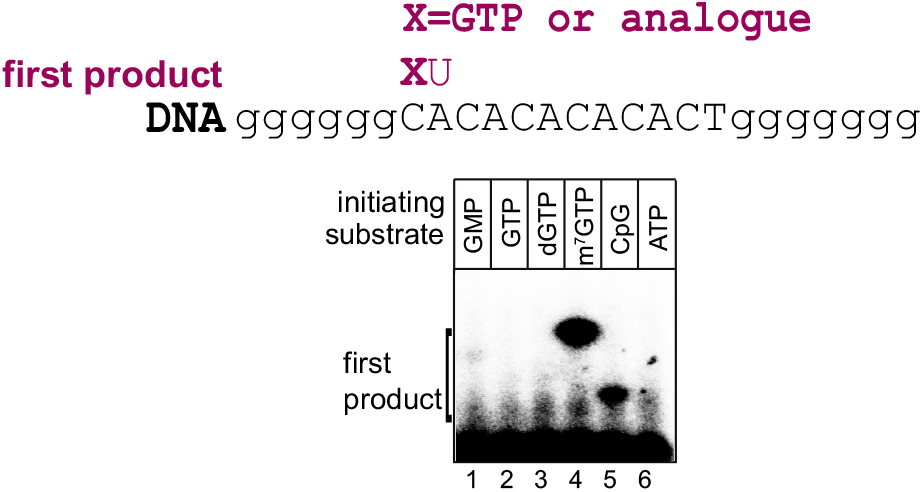
Human primase catalytic subunit P49 efficiently utilises m^7^GTP as initiating substrate. Scheme of the singlestranded DNA template is above the gel image. On the gel products of first step of synthesis with different initiating substrates and UTP as extending nucleotide are loaded.

## Discussion

Here we showed that DnaG primase initiates synthesis of a replication primer with ADP-containing cellular cofactors. This ability is reminiscent of that of RNA polymerases of transcription, yet there are notable differences. We found that DnaG uses NAD+ very inefficiently compared to NADH, in striking contrast to bacterial and mitochondrial transcriptases which do not distinguish between the two (4, 6). This feature of DnaG might connect priming of replication to the redox state of the cell. We also showed that the template base at −1 position does not play role in non-canonical substrate utilisation by DnaG, unlike transcriptases, which at least in some instances are sensitive to the −1 base (3). This result suggests that the incorporation of cofactors does not lead to spurious initiation or increased noise in the system.

For the first time here we show that global starvation response regulator, ppGpp affects non-canonical initiation of RNA synthesis. It remains to be seen if incorporation of non-canonical substrates by bacterial RNAP is influenced by ppGpp.

RNA pieces of Okazaki fragments are destined to be removed. Yet despite the transient nature of these RNAs, a balance between kinetics of removal and extension defines the mean size and length distribution of Okazaki fragments and ultimately replication kinetics (9). We showed that the rate of processing of replication primer by PolI is affected specifically by 5’-cofactors. NADH greatly stimulates, and FAD and DP-CoA inhibit the processing. We also found that decapping nuclease NudC can remove 5’-cofactors from RNA even if RNA is annealed to DNA. Therefore, hypothetically, NudC may assist removal of primers aberrantly left unprocessed in the cell.

The propensity of the human primase catalytic subunit to incorporate efficiently a modified substrate, an analogue of the classic cap m^7^GTP, suggests the probability of non-canonical initiating of replication in eukaryotes.

The evidence on physiological roles and the regulatory mechanisms of cofactors as non-canonical initiating substrates in transcription are emerging. It was demonstrated that capping influences the stability of RNA (2), and capping stimulates RNAP escape from the promoter (3). We show that replication may also be affected by cofactor incorporation. Currently, more potential nucleotide analogues, including dinucleotide polyphosphates incorporated into 5’ position of cellular RNAs are being discovered in both kingdoms (2). The role of these emerging substrates potentially extends to replication regulation.

Our results strongly suggest that cofactor initiation of replication happens *in vivo*, and future research would confirm this. At present we were unable to detect presence of cofactors on RNA primers *in vivo*, most probably due to the transient nature of a replication primer, relatively low number of primers per cell, and the sensitivity limitation of the methods we used. Nevertheless, we keep trying.

## Materials and Methods

### Strains and Plasmids

DnaG wt was overexpresed grom pCA24N-dnaG plasmid (KAYO collection), DnaB wt from pCA24N-dnaB (strain JW4012) ASKA collection. DnaG was transferred to plasmid pET28a via Gibson Assembly. DnaG mutants K229A, Y230A, F238A, K241A and D309A were generated by Quick change mutagenesis from pET28a-dnaG wt basis.

### Media and selection

*E. coli* were always grown in Luria bertani (LB) medium (broth or agar). In presence of pET28a Kanamycin (50 mg/L) was given to select for plasmid, pCA24N was selected with Chloramphenicol (34 mg/L).

### Cloning of WT and mutant DnaG

Gibson Assembly and QuickChange protocolls were used according to manufacturer’s specifications and transformed into *E. coli* DH5α cells. After sequencing, the plasmid was retransformed into *E. coli* T7 overexpression strain.

### Protein purification

DnaG wt and mutants were grown in 1 L LB with Kanamycin to an OD600= 0.5 at 37° C before induction with IPTG (1 mM) and continued growth at room temperature overnight. Cells were pelleted and stored at −80° C until lysis using sonication in Grinding Buffer (20 mM Tris-HCl pH 7.9, 200 mM NaCl, 5% Glycerol) and Ni-column purification (His-trap) in Purification Buffers A (20 mM Tris-HCl pH8, 600 mM NaCl, 5% g;ycerol) and B (same as A plus 200 mM imidazole). Protein was bound to column at 10 mM Imidazole, washed at 25 mM, and eluted at 200 mM imidazole). The reasonably clean protein was dialysed against Storage buffer (20 mM Tris-HCl pH 8, 50 % glyceerol, 200 mM KCl, 0.5 mM EDTA, 1mM DTT). DnaB was overexpressed from pCN24A in presence of Chloramphenicol, but likewise induced with 1 mM IPTG administered at OD600= 0.8 and grown over night. The rest was done as with DnaG. NudC protein was purified similarly except in the absence of EDTA and DTT.

### *In vitro* primer synthesis

Primers were synthesises by DnaG (+/−DnaB) on one of above mentioned templates in presence of 100 uM UTP and 10 uM GTP as well as approx 0.04 MBq αP32-GTP per uL in primase buffer (50 mM HEPES pH 7.0, 20 mM Mg-Acetate, 100 mM K-Glutamate, 10 mM DTT) at 30° C for 10 minutes, unless otherwise indicated in results. The reaction was stopped either in Stop buffer (1x TBE, 7 M Urea, 20 mM EDTA, 100 ug/ml heparin, 0.02% Bromphenol blue, 0.02 % Xylene cyanole, 85% Formamide) or by addition of 1 M NaCl (final concentration). Buffer exchange or removal of excess NTPs and abortive products were performed by gel filtration (BioRad Micro-Spin 6), if necessary.

### Pol I primer degradation

Primer was generated on the hairpin template (TTTACGCTTCGTTGACACACACACTGCGCGTTTGGGAAAACTCTTTCCCAAAC) by DnaG (1 μM) and DnaB (3 μM) premixed at RT with initiator (100 μM ATP or 1 mM other initiator (NAD, NADH, FAD, dpCoA)), 10 μM (α-^32^P) GTP (0.1 mCi/mmol), 100 μM UTP, in primase buffer (50 mM HEPES pH 7.5, 20 mM Mg-Acetate, 100 mM K-Glutamate, 10 mM DTT) at 30 °C for 10 min. Reaction was stopped by adding a final concentration of 1 M NaCl. Protein was removed from reaction by adding Ni-NTA agarose beads in primase buffer and 1 M NaCl. After incubation at RT for 5 min, bead supernatant was transferred to gel filtration column (Micro-Bio Spin 6, BioRad). Filtrate was added onto DNA Polymerase 1 (PolI) (*E. coli* source, Fisher Scientific), 0.25U/uL reaction, alongside 10 uM dNTPs, and PolI buffer (20 mM Tris-HCl pH 8, 50 mM KCl, 10 mM MgCl_2_). Incubation at 37°C was stopped after 5, 10, 20, 30, 45, 60, 120 and 180 seconds with Stop buffer.

